# Strigolactone-dependent gene regulation requires chromatin remodeling

**DOI:** 10.1101/2023.02.25.529999

**Authors:** J.L. Humphreys, C. Beveridge, M. Tanurdzic

## Abstract

Strigolactones (SL) function as plant hormones in control of multiple aspects of plant development. Regulation of gene expression by SL is a critical component of SL function. Immediate early gene regulation by SL remains unexplored due to difficulty in dissecting early from late gene expression responses to SL in whole plants. We used leaf-derived Arabidopsis protoplasts to explore early (5-180 minutes) changes in gene expression induced by SL by employing RNA-seq and ATAC-seq. We discovered over 1500 genes regulated by SL as early as 20 minutes, and up to 3669 genes across the entire time course of the experiment, indicative of rapid, dynamic regulation of gene expression in response to SLs. We identified 1447 regions of changing chromatin accessibility in response to SL that are likely to harbour SL cis-regulatory elements and cognate candidate trans-acting factors regulated early by SL. Importantly, we discovered that this extensive transcriptomic reprogramming requires the SYD-containing SWI/SNF chromatin remodelling complex(es) and regulates other chromatin remodellers. This study therefore provides the first evidence that SL signalling requires regulation of chromatin accessibility, and it identifies previously unknown transcriptional targets of strigolactones.

**One sentence summary:** Strigolactone regulated gene expression reprogramming requires chromatin remodelling by SPLAYED.

## Introduction

Plant development is shaped by plant hormones as the main regulatory signals that coordinate intrinsic developmental programs with environmental inputs^1,2^. They most often do so by engaging a variety of mechanisms to establish and maintain tightly regulated patterns of gene expression^3^. Transcriptional regulation depends on the activity of *trans*-acting transcription factors (TFs) at *cis* regulatory elements (CREs) in the genome. Accessibility of these genomic elements to the TFs is therefore critical for the regulation of gene function and is achieved through a dynamic process of chromatin remodelling, where upon perceiving a relevant signal, enzyme complexes can alter chromatin accessibility at specific locations throughout the genome^4–6^. Genetic studies have identified the role for chromatin remodelling in gibberellic acid (GA) ^7,8^, cytokinin (CK)^9,10^, auxin^11,12^ and abscisic acid (ABA)^13^, opening the way to genome-wide exploration of hormone signalling-related CREs^14,15^ including the most recently identified plant hormone, strigolactone.

The class of plant hormones called strigolactones (SL) act as a mediator of diverse aspects of plant growth and development such as axillary bud dormancy, leaf senescence and secondary growth^16–18^. In Arabidopsis, the direct targets of the SL receptor complex, such as the SUPPRESSOR OF MAX2-like 6, 7 and 8 (SMXL6/7/8) proteins are repressors of SL-response genes in the absence of SL^19,20^. This is achieved by an interaction between SMXL6/7/8 proteins with other transcription factors (TFs) and transcriptional co-repressors TOPLESS (TPL) and TPL-RELATED (TPR). SMXL6/7/8 proteins are degraded by the 26S proteasome system in response to SLs which would then lead to release of the early SL response from transcriptional repression.

Most investigations into transcriptome changes upon SL treatment have been performed following several hours/days of SL application, even though the rice SMXL6/7/8 orthologue D53 is degraded within ten minutes of SL application^21,22^. The complexity of tissues and organs involved in these experiments^23–26^ further complicates inferences in SL-mediated gene expression changes. Protoplasts (plant cells with the cell wall removed^27^) provide one solution to this confounding problem of sample complexity. Protoplasts are typically obtained by treatment of leaves or other plant parts with cellulase and pectinase enzymes that break down plant cell walls to produce spherical structures. Individual protoplasts in suspension provide a study system and have been used from a wide variety of species and sources to study hormone signalling and plant physiology^27–34^. Protoplasts can be quickly prepared from fully differentiated plant material and retain most of their physiological properties. as well as retaining cell type specific responses^35^. Arabidopsis leaves are the most common source of protoplasts since they are easy to harvest, high yielding, can be made from the large collection of available mutant lines and have capacity for transient expression assays^29^. Arabidopsis leaf protoplasts are able to respond to application of *rac*-GR24 treatments in a rapid and synchronised manner^36^.

Here, we use genome wide transcriptomic and chromatin profiling approaches combined with transient expression assays to infer the dynamics of SL-regulated gene expression and we provide the first evidence that SL signal leads to extensive changes in chromatin accessibility, most of which precede changes in expression of associated genes. We also identified the chromatin remodeller SPLAYED (SYD) is the key chromatin remodelling factor in SL signalling.

## RESULTS

### Transcriptional dynamics of SL signalling reveal key early gene expression regulation

To explore the early temporal dynamics of SL regulated gene expression, we established Arabidopsis leaf-derived protoplasts as an experimental system to study the transcriptional dynamics of SL responses on a time scale previously inaccessible using whole seedling treatments. We tested protoplast response to *rac*-GR24 by measuring gene expression of known targets of SL signalling. We found that *rac*-GR24 could induce SL-regulated TF *BRANCHED1* (*BRC1*)^37,38^ as well as a known target of *BRC1, HB21*^39^ within 20 and 45 minutes, respectively, and in a D14-dependent manner (Supplementary Figure 1). These results confirm that protoplasts are a suitable and robust system to study early SL-regulated gene expression.

To identify genome-wide gene expression changes in response to *rac*-GR24, we conducted RNA-seq analysis on *rac-*GR24 treated protoplasts at five time points (5, 20, 45, 90, and 180 minutes). We first identified differentially expressed genes (DEGs), that are regulated by *rac*-GR24 at any of the five time points, while removing any non-*rac*-GR24 effects on gene expression (Methods). In total, we identified 3769 DEG (Supplemental Table 1), regulated by *rac*-GR24 treatment at least at one time point. We found 1560 DEGs at 20 minutes, 108 at 45 minutes, 814 at 90 minutes and 2437 at 180 minutes (Figure 1A, Supplemental Table 1). No genes were found to be regulated by *rac*-GR24 at 5 minutes.

**Figure 1.**
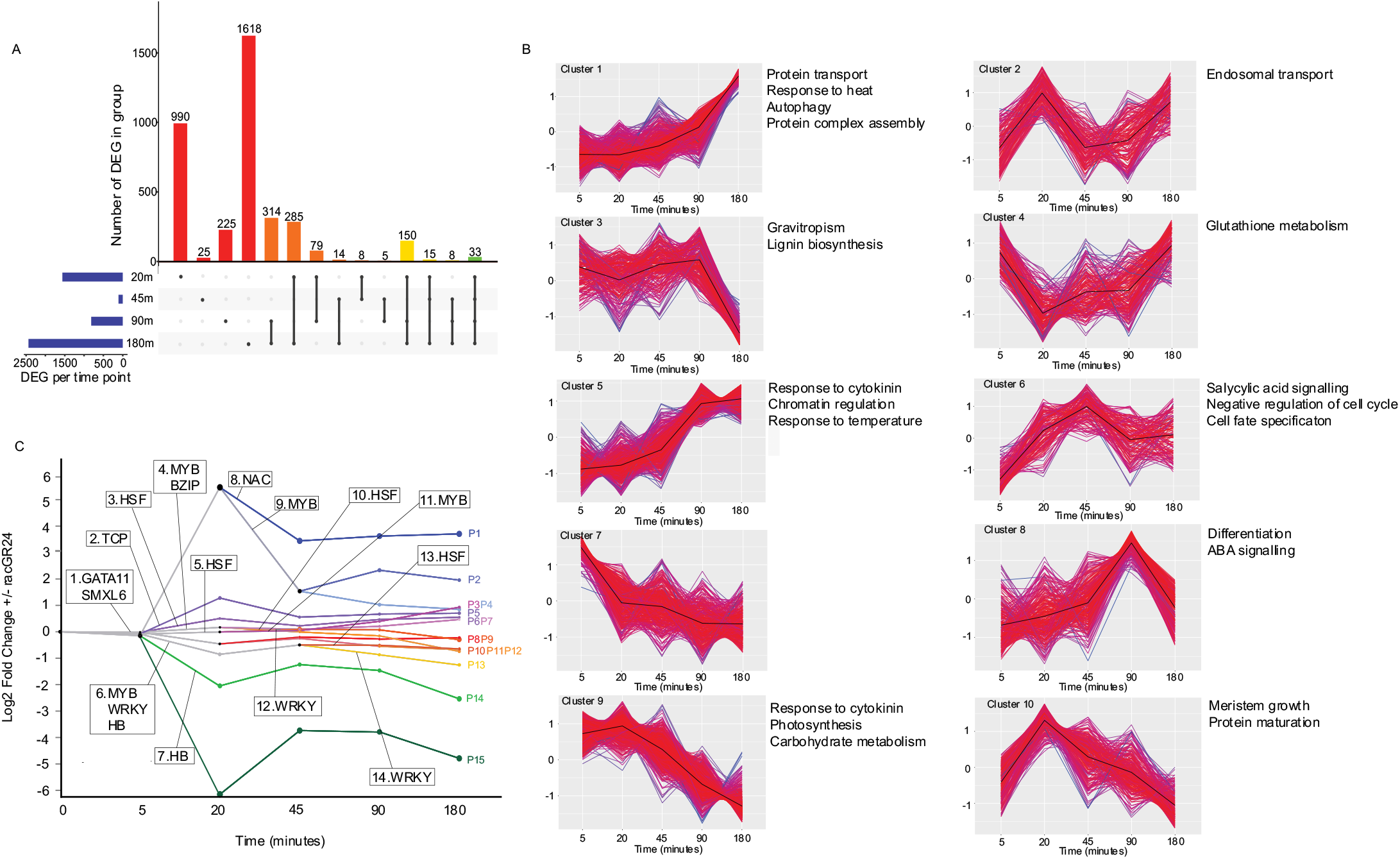
Dynamics of rapid gene expression changes in response to *rac*-GR24 in Arabidopsis protoplasts. **A)** Number of genes differentially expressed in *Arabidopsis* protoplasts after 20-, 45-, 90-, and 180-minutes of treatment with 1μM *rac*-GR24. Blue bars: total number of DEG at each time point, red bars: Number of DEG unique to each time point, orange bars: number of DEG common to two time points, yellow bars: number of DEG common to three time points, green bar: number of DEG common to all time points. Zero intersect values are not shown. **B)** k-means clustering of differentially expressed genes in response to 1uM *rac*-GR24 (5- to 180-minutes). Clusters are labelled C1-C10, and the top enriched GO terms in each cluster are shown to the right. **C)** DREM modelling of co-expressed genes in response to 1μM *rac*-GR24 with15 paths. The Y axis indicates the log2 fold change in expression in response to 1μM *rac*-GR24 and the X axis indicates the time in minutes. Black nodes indicate a split in a path. The TF families assigned to each DREM path are indicated.

We captured two waves of transcriptional responses to *rac*-GR24, the first at 20 minutes and the second starting at 90 minutes of *rac*-GR24 treatment. We predict that the first wave of transcriptional regulation in response to GR24 likely represents induction of early SL signalling containing transcriptional regulators. The second wave is indicative of downstream signalling cascades and biological processes induced or repressed by SLs. Importantly, we captured transient transcriptional responses to *rac*-GR24 supply; 63% of the genes regulated at 20 minutes were exclusively differentially expressed at that time point (Figure 1A). Our dataset, therefore, captures undescribed early and dynamic effects of SL signalling on transcriptional regulation and identifies novel early SL response genes.

We identified 342 TFs as SL-regulated, (Supplemental Table 2), with 138 of them differentially expressed by 20 minutes (with equal parts upregulated (67 (49%)) or downregulated (71 (51%)). We predict that the early transcriptional responses in SL signalling are at least in part modulated by activation or repression of these key upstream TFs. The TFs identified at 20 minutes include the TFs previously implicated in the SL response, *PAP1* and *BRC1* (Wang et al 2020), as well as novel SL-regulated factors such as: *GATA11, WRKY31, WRKY43, HSFA9*, and *BZIP24*.

Using k-means clustering to identify clusters of co-expressed genes in response to *rac*-GR24 we identified 10 clusters, labelled C1-C10 (Figure 1B, Supplemental Table 3). To explore the possible biological functions of SL-regulated DEGs in these clusters, we subjected our k-means clusters to Gene Ontology term enrichment analysis. The early responding/transient clusters (C2, C4, C6-8 and C10) showed little enrichment of GO term categories, while the late responding clusters (C1, C3, C5, C9) were enriched for multiple GO terms (Figure 1B). Increase in gene expression of genes involved in processes such as protein transport, autophagy (C1), negative regulation of cell cycle (C6), negative response to cytokinin, chromatin regulation and response to heat (C5) ABA signalling (C8), as well decrease in positive response to cytokinin, photosynthesis, and carbohydrate metabolism (C9) could be relevant to SL-regulated dormancy and senescence.

To further explore the regulatory cascade underlying dynamic SL transcriptional response, as afforded by our protoplast system, we implemented the Dynamic Regulatory Events Miner 2 (DREM2) algorithm^40^ to generate a temporal model of the SL-responsive gene-regulatory network (GRN). The DREM model identified 15 groups of co-expressed genes and predicted their transcriptional regulators (DREM paths named P1-P15 (Figure 1C). At the gene expression level, we identified a wide range of expression dynamics supporting previous k-means clustering results. Our model consists of seven upregulated paths (P1-7) and eight downregulated paths (P8-15, Figure 1C). Importantly, this analysis suggests that major regulatory events – indicated by bifurcations (splits) of paths - occur in the early wave of transcriptional responses to *rac*-GR24 (Figure 1C). This approach predicted numerous transcriptional drivers of changes in gene expression in the early time window, including SMXL6, and well-known TFs from the WRKYs, GATAs, REPRODUCTIVE MERISTEMS (REMs) and NO APICAL MERISTEM (NAMs) families. This is indicative of these TFs or members of the TF families regulating the first wave of differentially expressed genes at 20 minutes. At 20 minutes, bifurcations in paths begin to represent upregulated and downregulated genes. We identified members of the HSF, TCP, NAC, GATA and MYB TF families over-represented as regulators of the upregulated paths and members of the WRKY, HB, and BZIP families as regulators in the downregulated paths (Figure 1C). TCP, BZIP, HB and GATA factors were over-represented in the early paths and HSF, NAC, MYB, and WRKY factors were over-represented throughout the time series. The TFs are consistent with the TF families we identified as transcriptionally regulated at 20 minutes (Supplemental Table 2), and indicates a role for members of these families in driving gene expression responses to SL. Furthermore, the DREM2 model suggests that the SL response begins with a transient induction of SL response genes prior to sustained induction of late response genes, consistent with our k-means co-expression clusters (Figure 1).

### SL induces changes in chromatin accessibility ahead of transcriptional changes

Extensive transcriptional reprogramming in response to a signal is often accompanied by changes to chromatin accessibility. Supporting this, cluster C5 was found to contain several genes encoding chromatin regulators, such as SWI/SNF chromatin remodelling ATPase, BRAHMA (BRM) and other members of the SWI/SNF complex as well as the ISWI chromatin remodeller, PICKLE (PKL). We first explored whether chromatin remodelling is indeed required for SL-regulated gene expression. We tested gene expression in response to *rac*-GR24 in chromatin remodelling mutants *syd-5* and *brm-5*. SPLAYED (SYD) and BRM are the mutually exclusive ATPase components of the Switch2/Sucrose non-fermenting2 (SWI2/SNF2) chromatin remodelling complexes that have previously been implicated in transcriptional responses to plant hormone signalling^41^. We first identified that induction of *BRC1* expression in response to *rac*-GR24 requires SYD, as *BRC1* induction by *rac*-GR24 is lost in the *syd-5* mutant (Figure 2A). These results suggest that SYD acts upstream of *BRC1* in the strigolactone response pathway to regulate chromatin accessibility in order to permit expression of genes such as *BRC1*. On the other hand, we showed that the SYD paralog BRM is transcriptionally up-regulated by *rac*-GR24 5.7 fold only at 45 minutes but not sooner (Supplemental Table 1), implying that its response to *rac*-GR24 may require BRC1. Indeed, BRM up-regulation by r*ac*-GR24 is D14- and BRC1 -dependent (Figure 2B), and *BRM* is a direct target of BRC1, as determined by the TARGET assay^42^ (Figure 2B). Together, these results show that SYD activity is required for SL induction of *BRC1*, while *BRM* is directly regulated by BRC1, providing evidence that SWI/SNF chromatin remodellers BRM and SYD are required for SL-induced regulation of gene expression.

**Figure 2.**
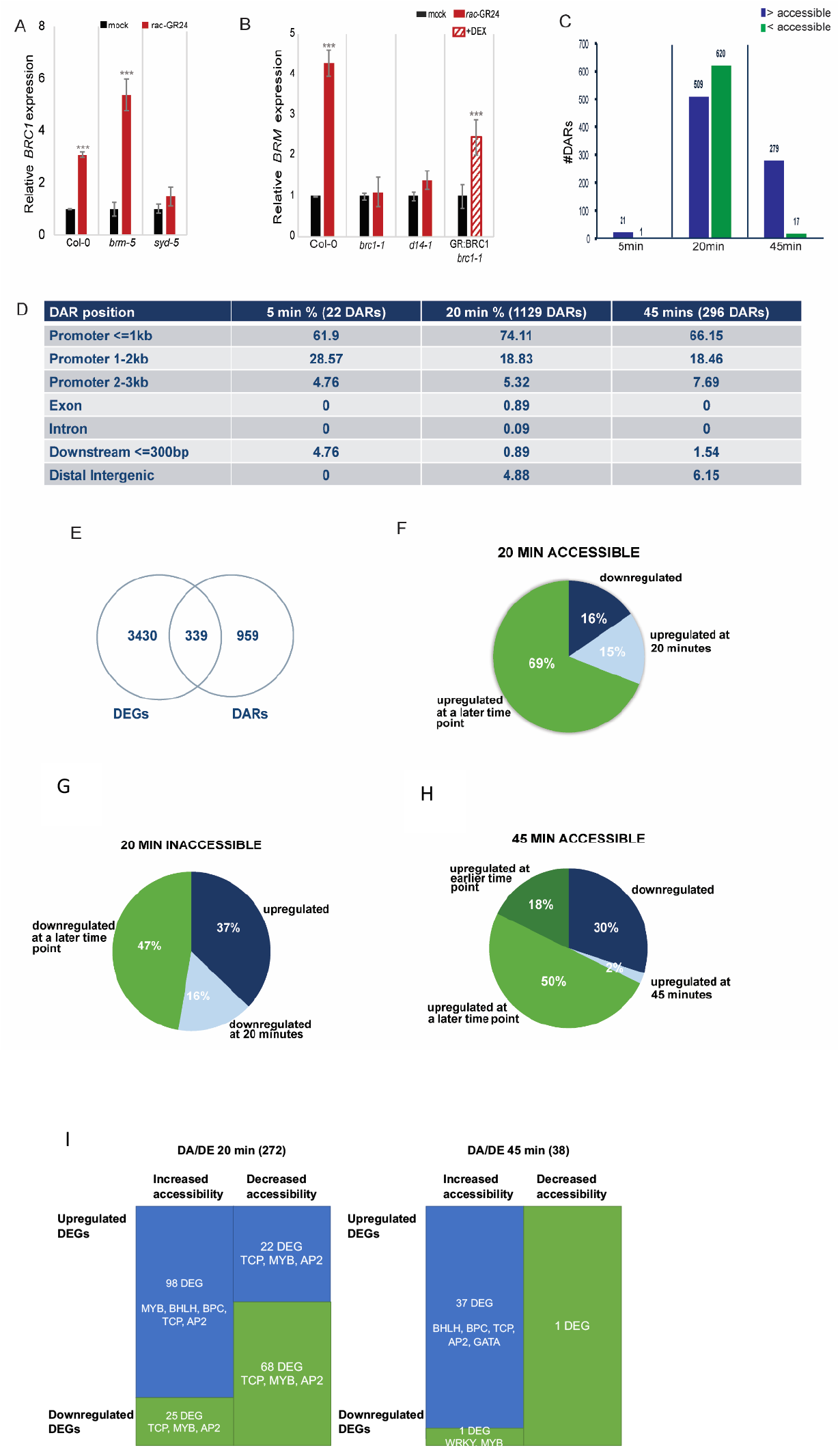
Chromatin accessibility dynamics in response to *rac*-GR24. **A)** Expression of *BRC1* in wild-type, *brm-5* or *syd-5*. Arabidopsis protoplasts treated with 1uM *rac*-GR24 or mock **B)** Expression of *BRM* in wild-type, *brc1-1, d14-1* or transfected *brc1-1* mutant cells. Arabidopsis protoplasts treated with 1uM *rac*-GR24 or mock (red bars). RHS bar with lines refers to induction of BRC1 protein into the nucleus by dexamethasone treatment. For A and B, Y axis represents relative gene expression means with error bars representing standard error n=3 independent biological replicates, (*P*<0.05 two-sided Student’s *t*-test). **C)** Total number of DARs found at each timepoint by DiffBind. >accessible: region increasing in accessibility, <inaccessible: region decreasing in accessibility, FDR <0.05. **D)** Genome distribution of annotated DA peaks for each time point. **E)** Venn diagram showing genes that are differentially expressed and differentially accessible (DA/DEs) upon treatment with 1uM rac-GR24 in WT for three biological replicates. **F-H)** Distribution DA/DEs with respect to the time of differential expression. **I)** Schematic representation of TF binding motifs enriched in DA/DEs at 20 minutes. Blue sections represent upregulated DE/DAs at each time point and green sections represent downregulated DA/DE’s at each time point.

### Genome-wide changes in chromatin accessibility in response to SL

To investigate the genome-wide targets of SL-dependent chromatin remodelling we performed the Assay for Transposase Accessible Chromatin with sequencing (ATAC-seq)^43^. Arabidopsis protoplasts were treated with 1 *μ* M or 0 *μ* M *rac*-GR24 for 5, 20 or 45 minutes. These early time points were chosen to complement the timing of changes in gene expression observed in the transcriptomic dataset, and to assess SL effects on chromatin accessibility before and after *BRC1* induction. Analysis of ATAC-seq data identified ~27,000 accessible chromatin peaks in each sample. We identified a total of 1447 Differentially Accessible Regions (DARs) in response to SL across the three time points: 22 DARs at 5 minutes, 1129 DARs at 20 minutes, and 296 DARs at 45 minutes (Figure 2C, Supplemental Table 4). Overall, of the 1447 DARs, 809 (55%) showed increased accessibility, whereas 638 (45%) were decreased in accessibility (Figure 2C). Just like the transcriptional response to SL, chromatin accessibility changes are also very dynamic; the majority of DARs at 5 and 45 mins increased in accessibility (Figure 2C) while slightly more regions were closed in response to SLs at 20 minutes (Figure 2C). The majority of DARs (65-75%) are found within 1kb of gene coding regions (Figure 2D), which may indicate that some of these regions may contain SL-responsive CREs. Each DAR was associated with its| putative target gene based on the nearest annotated transcriptional start site (TSS) using ChIPseeker^44^ (Supplemental Table 4). As multiple DARs can be annotated to one gene, 1298 unique genes were associated with the 1447 DARs (Supplemental Table 4).

To identify putative SL-responsive CRE within DARs and potentially associated TF families, we searched for DNA sequence motifs that were significantly enriched within the DAR sequences at each time point and compared them to the known TF-binding motifs in Arabidopsis^45^. The specific locations of these motifs and their associated TF within each DAR and its proximal gene were identified (Supplemental Table 5). Within DARs with increased accessibility in response to SL, the TF families with the highest number of TF binding motifs within DARs at 20 min were found for MYB, TCP and BPC TFs, while the TCP, BPC, bHLH and GATA TFBM were found within the DARs at 45 min (Supplemental Table 5). Within DARs with decreased accessibility, the TF families with the highest number of targets were TCP, AP2, bHLH, MYB, FRS and WRKY at 20 minutes, and MYB and WRKY at 45 minutes. A full list containing the number of DARs predicted to be targeted by each TF motif (p<1E-4) can be found in Supplemental Table 5. The TF DNA binding motifs identified in DARs are a subset of those identified as predicted regulators in the DREM model (Figure 1C). Furthermore, analyses of the overrepresented TF DNA binding motifs found in the promoters of DEGs in clusters (C1-C10) found at least one occurrence of these DNA binding motifs enriched in DEGs except the BPC family DNA binding motif. Taken together, these results suggest SL-regulated chromatin remodelling is required to make CRE’s accessible to TFs important for SL-regulated gene expression.

### SL mediated changes in chromatin accessibility precede changes in transcription

It is thought that dynamic regulation of chromatin accessibility leads to potential changes in gene expression by making CREs accessible or inaccessible to their corresponding TF ^46,47^.To determine the extent to which SL-dependent changes to chromatin accessibility correlate with changes in gene expression, genes closest to SL DARs (1298 genes) were compared with the DEG (3669 genes) from the transcriptome time series. This shows that 339 (32%) of the genes associated with changes in chromatin accessibility are differentially expressed in response to SL within the time frame of our experiments (Figure 3E). The 339 genes were labelled as differentially accessible and differentially expressed (DA/DE) (Supplemental Table 6). These observations mean that while chromatin remodelling is not required for all observed changes in gene expression in response to SL, a third of the SL-regulated transcriptome is associated with changes in chromatin accessibility, including MYB15, MYB30, MYB74, HSFB2B, WRKY42, WRKY47.

**Figure 3.**
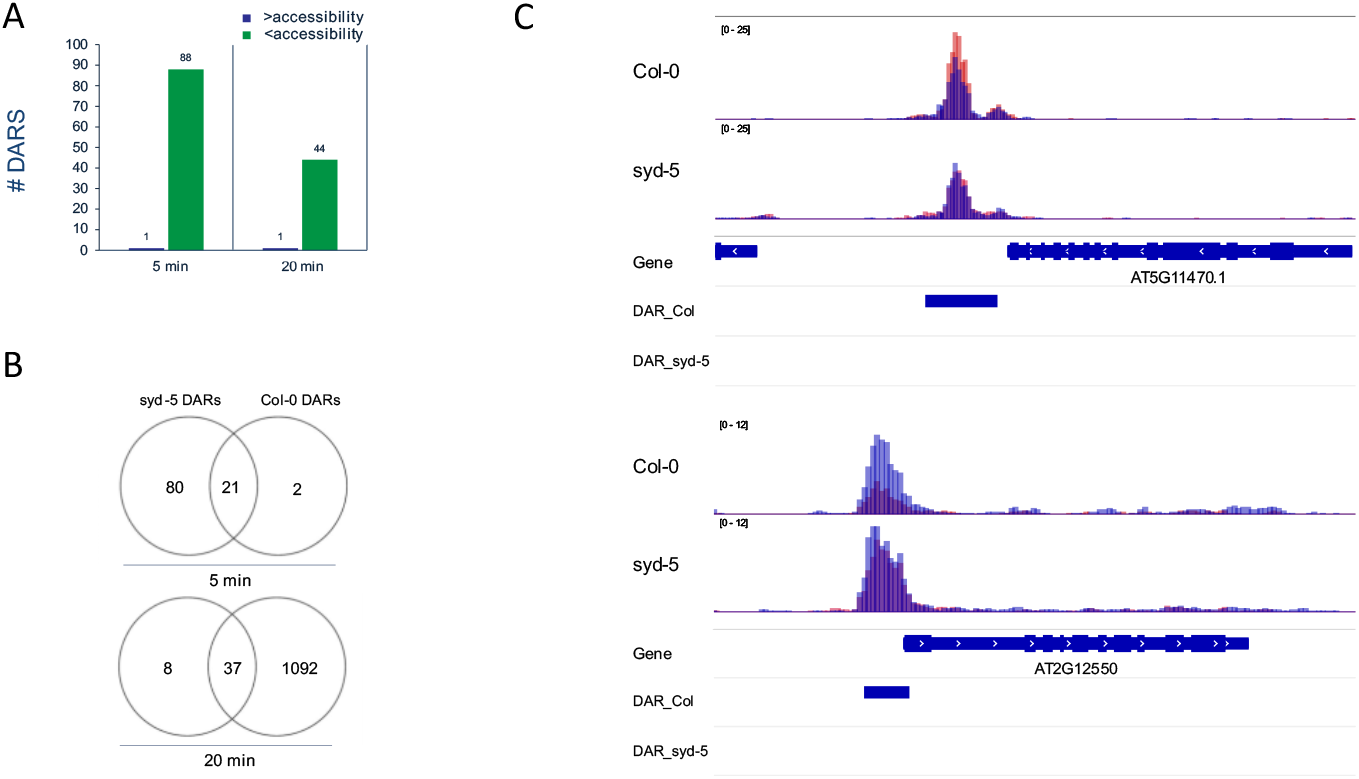
SL-regulated changes in chromatin accessibility require chromatin remodeller *SYD*. **A)** Total number of DARs found at each SL treatment timepoint in Col-0 (top) and *syd-5* (bottom). >accessible: region increasing in accessibility, <inaccessible: region decreasing in accessibility, FDR <0.05. **B)** Venn diagram showing DARs in common between WT and *syd-5* in response to 1uM *rac*-GR24. **C.)** Genome browser views of merged data sets showing changes in chromatin accessibility in response to *rac*-GR24 in Col-0 (top) but not *syd-5* (bottom). Blue bars indicate DARs identified by DiffBind. +/- *rac*-GR24 regions are overlaid, Blue: -*rac*-GR24 Red: +*rac*-GR24.

To further dissect the timing of SL-regulated chromatin accessibility changes on early changes in gene expression, we identified genes that show corresponding DA/DE changes at the same timepoint, as well as those where DA change precedes DE (Figure 2F-H). The majority of DA/DE genes become differentially expressed at a time point that is later than the change in chromatin accessibility, which suggests that SL-dependent changes in chromatin accessibility prime regulatory regions of DNA, rather than being an outcome of otherwise regulated transcription. By searching for occurrences of DNA binding motifs found enriched in DARs we were able to delineate TFs that were potentially regulating early vs later changes in DA/DE gene expression. For accessible DA/DE at 20min, the same regulators were identified for early vs late gene expression changes (Figure 2I). Contrastingly, at 45 minutes, GATA family TF binding motifs were exclusively found in 45 minutes DA/DE genes that were differentially expressed later in the time series (Figure 2I). For inaccessible DA/DE at 20min, GATA, BHLH and AP2 family motifs were exclusively found in genes that were regulated transcriptionally at later time points. These results suggest that GATA, AP2, and BHLHs TFs are required to regulate the late SL-regulated DA/DEs.

Collectively, the RNA-seq and ATAC-seq data uncover key early regulatory events occurring in response to SL. We highlight TFs that are regulated transcriptionally by SLs and are predicted to bind to SL-regulated DARs to regulate the expression of the genes adjacent to the DAR. A good example of this is *GATA11* which is transcriptionally induced by SLs at 20 minutes. *GATA11* is predicted to be an important driver of early gene expression events (Figure 1C, Supplemental Table 1) and is predicted to bind to SL-regulated DARs. Similarly, we identify eight SL-regulated WRKY TFs enriched in clusters C9 and C3 and DARs of downregulated DA/DE genes. Of these, *WRKY47* and *WRKY51* are transcriptionally induced by SLs at 20min. The observation that WRKY family DNA binding motifs are found enriched in late responding, downregulated clusters C3 and C9 suggests that induction of WRKY factors *WRKY47, WRKY51* (20 min) and *WRKY9* (45 min), leads to downregulation of SL-response genes. Therefore, this data supports the notion that chromatin accessibility changes are required for SL-mediated changes in gene expression and suggests that TCP transcription factors and other co-regulatory transcription factor families such as GATA and WRKY may be involved in modulating the SL-responsive transcriptome.

### Role of SPLAYED in strigolactone-induced changes in chromatin accessibility

As SYD is required for transcriptional induction of *BRC1* (Figure 2A), we explored the relationship between SYD and SL-mediated changes in chromatin accessibility. We performed an ATAC-seq on the *syd-5* mutant 5 minutes and 20 minutes after SL treatment. These time points were chosen as SYD is important for induction of *BRC1* expression by 20 minutes (Figure 2A). In this analysis, 98 DARs were identified at 5 minutes, 84 becoming more accessible and 14 became less accessible. At 20 minutes only 44 DARs were identified, 43 with increased accessibility and 1 with reduced accessibility (Figure 3B). When compared to the WT ATAC-seq results, we found that of the 1129 DARs found in WT at 20 minutes, 1092 (97%) were not differentially accessible in the *syd-5* mutant, and of the 22 DARs found at 5 minutes, 2 were not differentially accessible in the *syd-5* mutant (Figure 3A). This suggests that 97% of SL-regulated DARs at 20 minutes require chromatin remodelling protein SYD, reinforcing that SYD is required for the SL chromatin and transcriptional response. We found a large overlap between published SYD ChIP peaks^48,49^ and SL-regulated DARs. 47% of the SL-regulated DARs were also found to be targets of SYD. Interestingly, the majority of DARs identified at 5 minutes did not require SYD, indicative of a different factor regulating the 5-minute response.

## DISCUSSION

Strigolactones regulate many aspects of plant physiology and development similar to, and often in concert with other plant hormones like auxins or cytokinins ^18,50^. Previous studies using whole seedling treatment with SL over periods of time much longer than the time needed to activate degradation SMXL6/7/8 proteins identified a small number of differentially expressed genes ^23–26,51^ in comparison with other plant hormones ^52,53^. In this work we explored the immediate/early gene expression changes in response to SL by harnessing the power of leaf-derived protoplasts as an experimental system that enables measurement of early gene expression dynamics in a synchronized population of plant cells. We showed that expression of the key SL-activated transcription factor *BRC1* and its previously identified direct target *HB21* could be induced by 20- and 45-minutes, respectively, which persisted throughout the 3h SL treatment (Supplemental Figure 1). Detecting the earliest transcriptional changes in response to SL in this manner we identified early and late SL-response genes over a 3-hour period, with 1560 DEG in the first wave at 20 minutes (but not at 5 minutes) post SL addition and with 2441 DEG by 180 minutes (Figure 1A, Supplemental Table 1). This is consistent with the timing of SMXL6/7/8 degradation thought to occur within 10 minutes of SL application^21,22^. This result demonstrates that the protoplast system can facilitate the identification of thousands of genes at time points that have previously not been implicated in SL function. Importantly, the early wave of SL-regulated genes did not consist of as many enriched GO terms as the late wave (Figure 1B). This could be indicative of these genes being upstream or required for the second wave of SL-regulated gene expression (Figure 2B), a hypothesis supported by our dynamic modelling of the SL gene regulatory network, which included SMXL6, BRC1/TCP, GATA11 and WRKY family transcription factors (Figure 1C). These results vastly increase the number of SL-regulated genes discovered in Arabidopsis and identify novel candidate regulators of SL signalling in Arabidopsis cells.

### Chromatin remodelling is a component of strigolactone signalling

Chromatin is a dynamic structure that responds to stimuli by changes in its accessibility to trans-acting regulators, or by changes to its covalent modifications of DNA and histone molecules. This ultimately play key roles in regulating the transcriptional output. To determine whether chromatin regulation is a part of SL signalling, we profiled chromatin accessibility in response to SL using ATAC-seq and integrated this dataset with the RNA-seq dataset. We demonstrate that SL signal changes chromatin accessibility at 1447 locations in the Arabidopsis genome, with earliest changes occurring as early as 5 minutes. The greatest changes in chromatin accessibility occurring 20 minutes after SL-treatment (Figure 2B). This is consistent with a large change in gene expression observed at 20 minutes (Figure 1A). We further showed that the majority of SL DARs are found in gene promoters, consistent with similar experiments looking at chromatin accessibility changes in response to hormones, temperature and light changes in Arabidopsis ^14,54,55^. Importantly, we were able to identify the TF binding motifs within the 20 minute-accessible-DARs matching those of the TF families regulated by SLs at the early time points (Figure 1C, Supplemental Table 2). These results further corroborate gene regulatory network model (Figure 1C that predicts SMXL6 TCP, MYB, AP2, WRKY and GATA family members as important regulators of gene expression. Our results also show for the first time that SL signal requires regulation of chromatin accessibility at the *BRC1* locus (Figure 2A), likely in the presumptive cis-regulatory regions of *BRC1* that also contains the SMXL6, MYB and MADS box domain TF binding motifs and DNA motifs found enriched in SYD ChIP-seq peaks^48^ (Supplemental Figure 2).

Our results identify chromatin remodelling as a likely critical component of SL signaling, as has been already been shown in other plant hormone signalling pathways^11,13,41,56^, SWI/SNF chromatin remodellers have been implicated in increasing accessibility at promoters of genes involved in cytokinin signalling^10^, auxin signalling^11^, and reducing accessibility of genes involved with GA signalling^8^, and ABA signalling^13^.

### Strigolactone regulated changes in chromatin accessibility precede transcriptional changes

SL-induced changes in chromatin accessibility that we identified comprised both increases and decreases in chromatin accessibility in response to SL, mirroring the directions in gene expression change (Figure 1). 32% of SL-regulated DARs occurred in or near genes whose expression was also altered by SLs (DA/DE genes, Figure 3E) showing either increased accessibility and induction of gene expression or decreased accessibility and repressed gene expression (Figure 3F-H). This is consistent with the current model of accessible chromatin being associated with increased gene expression and inaccessible chromatin associated with repression of gene expression^46,47,57^. Furthermore, for 54% of DA/DE genes, changes in accessibility precede changes in gene expression (Figure 3F-H). These results indicate that SL-dependent changes in chromatin accessibility likely identify some of the SL cis-regulatory regions of SL-regulated genes, which are further utilized when a TF-complex becomes available/expressed. This may suggest a SL-regulated mechanism of targeting genomic locations for changes in chromatin accessibility. It is thought that SMXL/TPL complex can play a role in inducing a repressive chromatin complex^11,58^. Degradation of SMXL6/7/8 proteins upon SL perception could therefore be is an important component of modulating accessibility cis-regulatory regions of SL-responsive genes. On the other hand, SL-induced decrease of chromatin accessibility at hundreds of regions in the genome would require induction and localization of a different repressive complex.

### Strigolactone regulated chromatin remodelling depends on the SWI/SNF chromatin remodellers SPLAYED (SYD) and BRAHMA (BRM)

Our initial observation that *BRM* is upregulated in response to SLs lead us to further investigate BRM and its homologue SYD as prime chromatin remodelling candidates for SL-initiated changes in chromatin accessibility and gene expression (Figure 2A). BRM and SYD both have strong pleiotropic phenotypes making it difficult to study their responses to specific hormones ^59–62^. Using *syd-5* mutant protoplasts, we showed that SYD is required for the induction of *BRC1* (Figure 2A), while BRC1 directly regulates *BRM* expression (Figure 2B), creating a hierarchy of SYD- and BRM-dependent regulation of gene expression in response to SL.

Using ATAC-seq in the *syd-5* mutant we were able to show that SYD is for 97% of SL DARs in wild type protoplasts (Figure 3). Importantly, evidence for physical localization of SYD-containing complexes to SL DARs is provided by previously published SYD ChIP-seq data^48,49^, with a 48% overlap between SL DARs and SYD ChIP-seq peak regions. These results place SYD as a critical chromatin remodeller early in the SL response and further implicates a SWI/SNF chromatin remodelling complex member in the SL response. Its coordinated targeting in response to SL could be accomplished by either specific interaction between the complex and a DNA binding factors specifically induced by SL, or via association of the SYD complex with genomic regions specifically marked for this recruitments via an unknown mark/mechanisms. An example of a targeted mechanism is the auxin-regulated chromatin switch^11^, where upon auxin sensing the MONOPTEROUS transcription factor recruits SWI/SNF chromatin remodelling ATPases (SYD or BRM) to increase accessibility at genes important for flower primordia initiation. An important question to be answered in the future is how non-sequence specific chromatin remodelling complexes containing SYD or BRM can function in a sequence- and function-specific context to change gene expression in a SL-dependent manner.

### Final conclusion

Transcriptional reprogramming often requires changes in chromatin accessibility state. Here, we identified that extensive transcriptional reprogramming by the plant hormone strigolactone is quick upon SL perception, and that this reprogramming is accompanied, and often preceded by changes in chromatin accessibility at genomic location likely to contain *cis*-regulatory regions of SL-regulated genes. Chromatin remodellers SYD and BRM are involved in SL-induced chromatin remodelling, and we provide first evidence the chromatin remodeller SYD is a critical component of SL signalling.

## METHODS

### Growth of Arabidopsis

All experiments using wild-type Arabidopsis thaliana were done using the Arabidopsis thaliana Columbia-0 ecotype (Col-0). Transgenic *d14-1* (CS913109), *brc1-1* (CS90653), *syd-5* (SALK_023209), and *brm-5* (CS68980). Arabidopsis lines used were in a Col-0 background. Arabidopsis plants were grown in soil mix supplemented with dolomite (1g/L) and osmocote 2g/L. Seeds were stratified for two days before being placed in a Thermoline growth cabinet at 22°C with a 16hr-light (150umol m/s2) /8hr-dark light regime. Plants between 3-4 weeks old before floral transition were selected for experiments.

### Protoplast Isolation and *rac*-GR24 treatments

In all protoplast experiments, rosette leaves were taken from 4-week-old plants prior to floral transition. One million cells were isolated per biological replicate for experiments. 10/12 Arabidopsis leaves yielded three million viable cells. The upper epidermal surface of leaves was stabilised to a piece of PVC electrical tape. The lower epidermis was fixed to another piece of tape. The lower epidermis was then carefully peeled away from the upper epidermis as described^32^. The peeled leaves with upper epidermis and mesophyll cells still adhered to the tape were transferred to a petri dish with 10mL of enzyme solution (1% cellulase, Onozuka R10 (Yakult, Tokyo, Japan), 0.2% macerozyme, ‘Onozuka’ R10 (Yakult), 0.4 M mannitol, 10 mM CaCl2, 20 mM KCl and 20 mM MES, pH 5.7). The leaves were shaken at 40rpm on a platform shaker at room temperature for 60 minutes. Released protoplasts were filtered through a 50 *μ* M CellTrics filter and centrifuged for 5 minutes at 100 x g. Protoplasts were washed in enzyme buffer (0.4 M mannitol, 10 mM CaCl2, 20 mM KCl and 20 mM MES, pH 5.7). Cells were counted using a hemocytometer and resuspended in MMg solution (0.4 M mannitol, 15 mM MgCl2, and 4 mM MES, pH 5.7) to a final concentration of 2 to 5 × 105 cells/mL. *rac*-GR24 treatment of protoplasts were conducted with three biological replicates (1 million cells per replicate). Protoplast suspensions were treated with 10 *μ* L of aqueous solution containing either 0 or 1 *μ* M of rac-GR24, in acetone. Protoplasts were collected consecutively at 5, 20, 45, 90 and 180 minutes following treatment by centrifugation (1000x g for 3minutes) and immediately frozen in liquid nitrogen for RNA-seq. RNA was extracted from samples using QIAGEN RNeasy mini kit as per the manufacturers instructions (catalogue number 74106). For ATAC-seq, 50,000 cells were collected consecutively at 5, 20 and 45 minutes and immediately processed for ATAC-seq library preparation.

### ATAC-seq library preparation

ATAC-seq library preparation protocol^63^ was modified for protoplasts. 50,000 treated protoplasts were immediately lysed in prechilled Nuclei Extraction Buffer 1, (1 ml; 15 mM Tris-HCl pH 7.5, 20 mM NaCl, 80 mM KCl, 0.5 mM spermine, 5 mM *β*-mercaptoethanol, 0.2% (v/v) TritonX-100, 1X Roche protease inhibitor tablets). The lysed cells were then filtered through 30 *μ* M CellTrics filters and centrifuged at 1200x g for 10 minutes. The pellet was resuspended in Nuclei Extraction Buffer 2 (1mL: 0.25M sucrose, 10mM Tris-HCL pH8, 10mM MgCL2, 1% (v/v) TritonX-100, 1X Roche Complete Protease Inhibitor) and centrifuged at 1200x g for 10minutes. The pellet was then resuspended in NEB3 (300 *μ* L: 1.7M sucrose, 10mM Tris-HCL pH8, 2mM MgCL2, 0.15% (v/v) TritonX-100, 1X Roche Complete Protease Inhibitor), then layered on top of 300 *μ* L NEB3 and centrifuged at 12,000x g for 10 minutes. The nuclei pellet was then immediately resuspended in 25 *μ* L of 2x tagmentation buffer containing 2 *μ* L of TDE1 (Illumina). The reaction was placed at 37C for 30 minutes with gentle mixing three times. Tagmented DNA was purified using a QIAGEN Minelute PCR purification kit and eluted in 11 *μ* L of elution buffer. Initially, 10 *μ* L of transposed DNA was used to undergo an initial 5 cycles of amplification using New England BioLabs Next High-Fidelity 2x PCR mastermix (M0541S) in 50 *μ* L reactions. To confirm the total cycle number needed for each sample, 5 *μ* L of the PCR reaction was used for RT-qPCR analysis. The cycle number at which the fluorescence for a given reaction at 1/3 of its maximum equals the additional number of cycles that each library required for adequate amplification. The remaining 45 *μ* L of PCR product was amplified for (N) cycles. Libraries were purified using AMPure beads, eluting in 22 *μ* L of elution buffer. Libraries were quantified using Qubit and pooled for Illumina HiSeq (PE 150).

### RNA-seq pre-data processing and differential gene expression analysis

Three libraries consisting of three biological replicates were prepared for each condition. A list describing each library can be found in Supplemental table 2-7. RNA was extracted using the Qiagen RNeasy mini kit. Total RNA was quantified using Qubit and 2ng of total RNA was sequenced at Novogene Co LTD. Libraries were sequenced using Illumina HiSeq 2500 and run on a 150 paired end cycle cartridge. Each library had over 25 million reads and quality control was carries out using fastQC^64^. Adapter sequences were removed as well as trimming and filtering the reads by Trimmomatix^65^. Read counts were generated for each transcript/gene by Salmon^66^. All libraries had >90% mapping percentage. Read counts generated by Salmon quantification were used as input for DESeq2. First, differential gene expression analysis was performed using DESeq2 version 1.18.1^67^. To identify DEG we used a design formula that takes into account the contrasts between two treatment levels (time and treatment) at FDR<0.05 (~time + treatment + time:treatment). At each time point, genes that were regulated by the interaction between SL treatment and length of treatment were determined based on the adjusted p-value for the significance of time:treatment interaction effect, using FDR multitesting error correction (p_adj_ <0.05). DESeq2 normalises and removes low counts internally. A complete list of DEG for each time point along with the foldchange and adjusted P values is available in Supplemental Table 1. Venn diagrams comparing gene sets were generating using the UpSetR package^68^.

### K-means clustering

Reads were normalised using counts per million (CPM) to normalise the depth effect of each library. CPM normalised expression matrix was scaled to a Z-scale matrix and the k-means function in Rstudio was used to cluster genes with the following parameters (set.seed(200000), centres = 10, iter.max=30). Plots were generated with ggplot2, dplyr, tidyr, and pheatmap.

### GO term overrepresentation

GO term enrichment was performed using the clueGO cystoscope plugin^69^, identifying enriched GO terms with adjusted p-values of <0.05. A minimum of 3 genes per cluster were required for enrichment.

### DREM2 analysis

The DREM model was generated using DREM v2.0.3^40^. Three files were used as input: A file listing the 33,323 TAIR10 gene IDs, a file containing previously identified gene-TF interactions and a file containing the log2FC of the DEG from the RNA-seq experiments. The TF interaction file that includes gene-TF interactions from AGRIS and DAP-seq was adapted^70^, to include gene-TF data of BRC1 and SMXL6. The expression data file contains the log2 expression values obtained from DESeq2 output for the 3769 DEG. NA values were replaced with blanks. Using the described input files, DREM models were generated without using the gene-TF interaction data and used the Train-test framework. DREM2 groups genes with similar expression patterns together and assigns putative TFs to gene sets based on previously determined TF-gene interactions ^40^. DREM2 enables the integration of time series gene expression data with static interaction data to predict time-point specific transcriptional drivers. To construct the DREM2 model, we input the log2FC gene expression values from the interaction model (~Time+Treatment+Time:Treatment) of the 3769 genes (FDR < 0.05) from our time course DEG analysis. The DREM model was generated based on our expression data alone and TF predictions were subsequently determined as in ^70^. This ensures groups of genes are assigned together based on their expression data rather than the mixture of *in vivo* and *in vitro* derived interactions from the TF-interaction file.

### ATAC-seq data analysis

Reads from RNA-seq and ATAC-seq libraries were trimmed based on quality using trimmomatic^65^. Reads were mapped to the Arabidopsis genome (Tair10) using Bowtie2^71^, keeping fragments with lengths up to 1000bp and allowing for 2 mismatches. The resulting BAM file was sorted and processed to remove duplicate and organelle reads. For IGV analysis and image capture, biological replicates were using MergeBam in Picard tools and converted BAM to bigwig using the bamCoverage function within deepTools^72^, with a bin size of 10bp and normalising by CPM. To call peaks for differential accessible region analysis MACS2^73^ was used. SPOT scores were >0.5 for all libraries, FRiP scores were >0.5 for all libraries. Differential accessible regions were identified using DiffBind^74^ with DESeq2 with the parameter “min members = 2”. The differential regions were annotated using the ChIPSeeker v.1.12.133 package^44^ within R using the annotatePeak function with promoters defined as –3000 to 3000bp from the transcription start site.

### Motif Enrichment

STREME^75^ from the MEME suite was used to identify motifs enriched in the promoters of genes present in individual clusters. All motifs were then compared with DAPseq motifs^45^ using TOMTOM^76^ to search for motifs overlapping by at least 5bp. FIMO^77^ was used to scan promoter sequences for candidate binding sites for motifs of interest.

### Quantitative Real Time PCR analysis

Reverse transcription of 500 ng of RNA was performed using the iScript reverse transcription kit (Bio-Rad) as per the manufacturer’s instructions. Real-time PCR was performed using SYBR green supermix (Bioline) as per the manufacturer’s instructions on a CFX384 Touch real-time PCR detection system. A melt curve analysis was included for quality assurance. The real-time data was processed in CFX Manager 2.1 software and LinRegPCR (Academic Medical Centre, Amsterdam, The Netherlands) and Microsoft Excel as previously described (Dun et al., 2012). Gene expression was normalized against both a geomean of reference gene expression *ELONGATION FACTOR 1 ALPHA* and the expression in the control samples. A list of primers used for these experiments can be found in Supplemental Table 8.

### TARGET induction of BRC1

TARGET^42^ experiments were conducted with 3 biological replicates with 10^6^ cells per replicate. *brc1-1* protoplast suspensions were transfected with a polyethylene glycol treatment using 120 *μ* g of plasmid. Protoplast suspensions were incubated at room temperature overnight while shaking at 40rpm on a platform shaker. The following morning, protoplast suspensions were pre-treated with 35 *μ* M cycloheximide (CHX; Sigma-Alrich) for 30minutes, after which either 0 or 10 *μ* M of dexamethasone (DEX, Sigma-Alrich) was added and cells were incubated at room temperature for 2hours. Treated protoplast suspensions were sorted with a BD Influx (BD biosciences) using 488nm excitation and measuring emission for red fluorescence at 610/20nm for red fluorescence. Cells were directly sorted into RNA extraction buffer and immediately processed using QIAGEN RNeasy mini kit.

## DATA AVAILABILITY

Data will be available through SRA accession#

## Acknowledgements

This work was supported by the Australian Research Council grant DP150102086 to CB and MT.

---

**Supplemental Figure 1.**
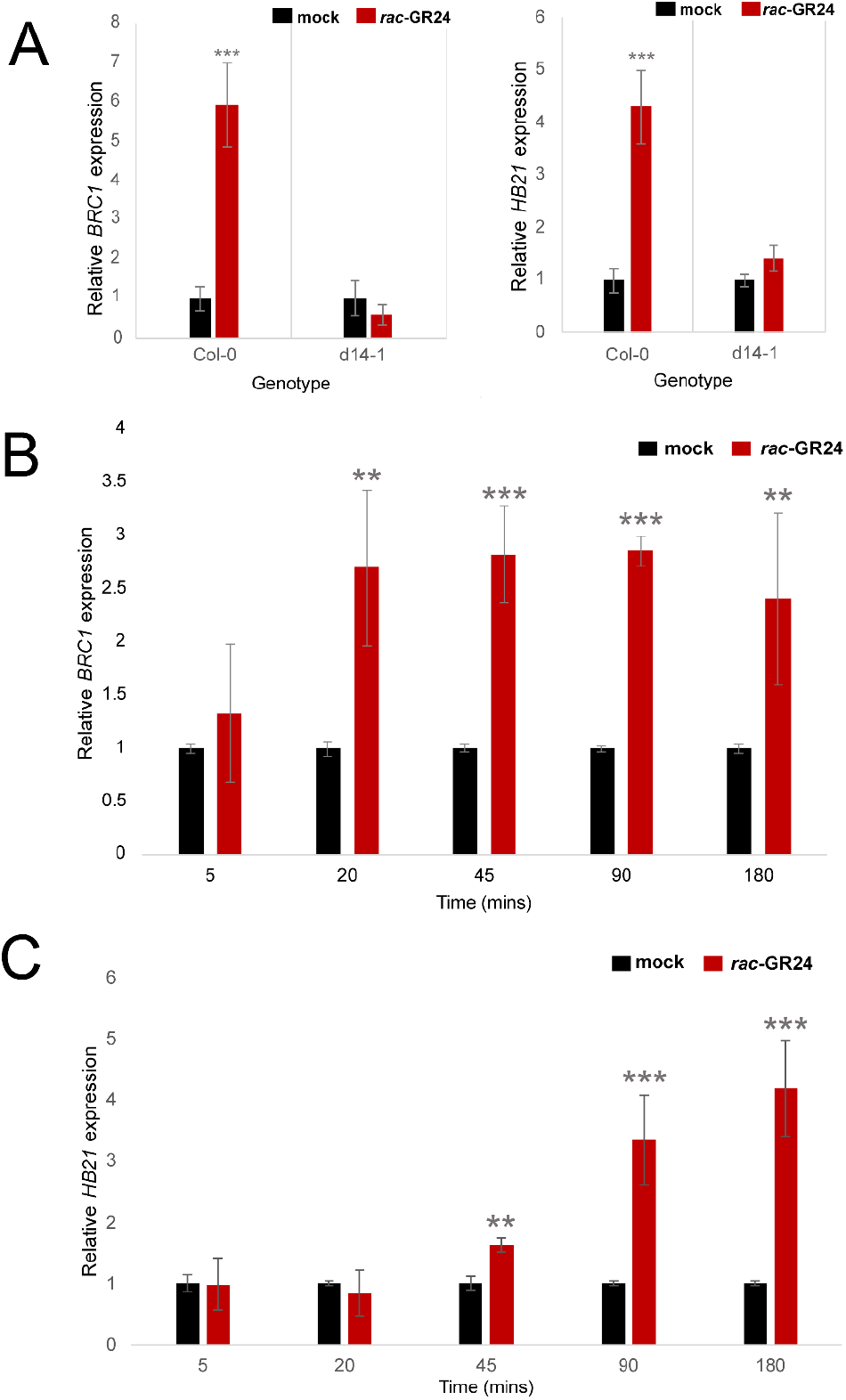
GR24 induces known transcriptional responses in Arabidopsis protoplasts. **A)** Relative expression of *BRC1* and *HB21* in wild-type (Col-0) and *d14-1* mutant Arabidopsis protoplasts in response to mock or 1μM *rac*-GR24 treatment determined by qRT-PCR. **B)** Relative expression of *BRC1* in Col-0 Arabidopsis protoplasts in response to mock or 1 μM *rac*-GR24 treatment for 5, 20, 45, 90, or 180 minutes. **C)** Relative expression of *HB21* in Col-0 Arabidopsis protoplasts in response to mock or 1μM *rac*-GR24 treatment for 5, 20, 45, 90, or 180 minutes. All data are means +/- SE, n=3 biologically independent replicates. **<0.01, ***<0.001

**Supplemental Figure 2.**
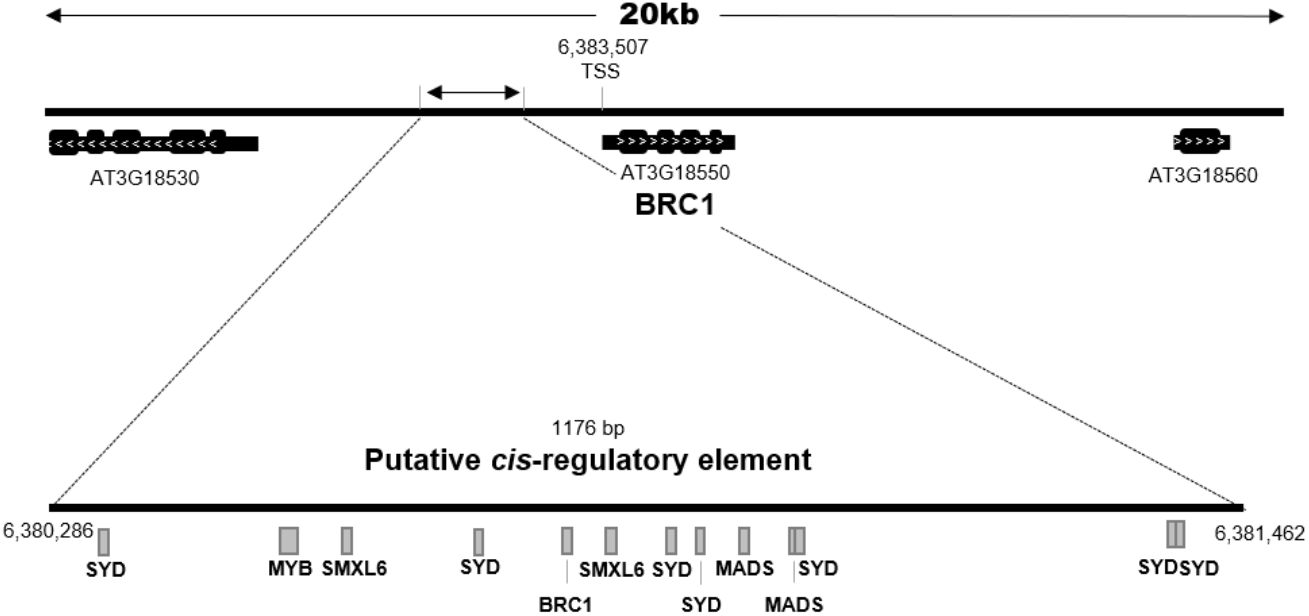
*BRC1* is a target of SL-regulated changes in chromatin accessibility. A Schematic representation of Genome browser views ATAC-seq datasets showing SL-induced chromatin accessibility changes occurring near *BRC1* with a schematic representation of the *BRC1*-DAR showing motifs found within the sequence. *CRE* was identified as a DAR using the Likelihood Ratio Test (LRT) implemented in DESeq2. Region is annotated by genome co-ordinates from Araport 11 (Tair 10).

## References

1. Benkova, E. Plant hormones in interactions with the environment. Plant Molecular Biology 91, 597–597 (2016).

2. Vanstraelen, M. & Benková, E. Hormonal Interactions in the Regulation of Plant Development. Annual Review of Cell and Developmental Biology 28, 463–487 (2012).

3. Blázquez, M. A., Nelson, D. C. & Weijers, D. Evolution of Plant Hormone Response Pathways. Annual Review of Plant Biology 71, 327–353 (2020).

4. Cosgrove, M. S., Boeke, J. D. & Wolberger, C. Regulated nucleosome mobility and the histone code. Nat Struct Mol Biol 11, 1037–1043 (2004).

5. Clapier, C. R., Iwasa, J., Cairns, B. R. & Peterson, C. L. Mechanisms of action and regulation of ATP-dependent chromatin-remodelling complexes. Nature Reviews Molecular Cell Biology 18, 407–422 (2017).

6. Sundaramoorthy, R. Nucleosome remodelling: structural insights into ATP-dependent remodelling enzymes. Essays Biochem 63, 45–58 (2019).

7. Ogas, J., Kaufmann, S., Henderson, J. & Somerville, C. PICKLE is a CHD3 chromatin-remodeling factor that regulates the transition from embryonic to vegetative development in Arabidopsis. Proceedings of the National Academy of Sciences 96, 13839–13844 (1999).

8. Archacki, R. et al. BRAHMA ATPase of the SWI/SNF Chromatin Remodeling Complex Acts as a Positive Regulator of Gibberellin-Mediated Responses in Arabidopsis. Plos One 8, (2013).

9. Furuta, K. et al. The CKH2/PKL Chromatin Remodeling Factor Negatively Regulates Cytokinin Responses in Arabidopsis Calli. Plant and Cell Physiology 52, 618–628 (2011).

10. Efroni, I. et al. Regulation of Leaf Maturation by Chromatin-Mediated Modulation of Cytokinin Responses. Developmental Cell 24, 438–445 (2013).

11. Wu, M. F. et al. Auxin-regulated chromatin switch directs acquisition of flower primordium founder fate. Elife 4, (2015).

12. Chung, Y. et al. Auxin Response Factors promote organogenesis by chromatin-mediated repression of the pluripotency gene SHOOTMERISTEMLESS. Nat Commun 10, 886 (2019).

13. Peirats-Llobet, M. et al. A Direct Link between Abscisic Acid Sensing and the Chromatin-Remodeling ATPase BRAHMA via Core ABA Signaling Pathway Components. Molecular Plant 9, 136–147 (2016).

14. Potter, K. C., Wang, J., Schaller, G. E. & Kieber, J. J. Cytokinin modulates context-dependent chromatin accessibility through the type-B response regulators. Nature Plants 4, 1102 (2018).

15. Wu, L.-Y. et al. Dynamic chromatin state profiling reveals regulatory roles of auxin and cytokinin in shoot regeneration. Developmental Cell 57, 526–542.e7 (2022).

16. Gomez-Roldan, V. et al. Strigolactone inhibition of shoot branching. Nature 455, 189–U22 (2008).

17. Umehara, M., Hanada, A., Magome, H., Takeda-Kamiya, N. & Yamaguchi, S. Contribution of Strigolactones to the Inhibition of Tiller Bud Outgrowth under Phosphate Deficiency in Rice. Plant and Cell Physiology 51, 1118–1126 (2010).

18. Waters, M. T., Gutjahr, C., Bennett, T. & Nelson, D. C. Strigolactone Signaling and Evolution. Annual Review of Plant Biology 68, 291–322 (2017).

19. Wang, L. et al. Strigolactone Signaling in Arabidopsis Regulates Shoot Development by Targeting D53-Like SMXL Repressor Proteins for Ubiquitination and Degradation. Plant Cell 27, 3128–3142 (2015).

20. Soundappan, I. et al. SMAX1-LIKE/D53 Family Members Enable Distinct MAX2-Dependent Responses to Strigolactones and Karrikins in Arabidopsis. The Plant Cell 27, 3143–3159 (2015).

21. Jiang, L. et al. DWARF 53 acts as a repressor of strigolactone signalling in rice. Nature 504, 401–405 (2013).

22. Patil, S. B. et al. Sucrose represses the expression of the strigolactone signalling gene D3/RMS4/MAX2 to promote tillering. http://biorxiv.org/lookup/doi/10.1101/2020.11.10.377549 (2020) doi:10.1101/2020.11.10.377549.

23. Lantzouni, O., Klermund, C. & Schwechheimer, C. Largely additive effects of gibberellin and strigolactone on gene expression in Arabidopsis thaliana seedlings. Plant J. 92, 924–938 (2017).

24. Wang, L. et al. Transcriptional regulation of strigolactone signalling in Arabidopsis. Nature 583, 277–281 (2020).

25. Min, Z. et al. Transcriptome analysis revealed hormone signaling response of grapevine buds to strigolactones. Scientia Horticulturae 283, 109936 (2021).

26. Ravazzolo, L. et al. Strigolactones And Auxin Cooperate To Regulate Maize Root Development and Response to Nitrate. Plant and Cell Physiology pcab014 (2021) doi:10.1093/pcp/pcab014.

27. Confraria, A. & Baena-González, E. Using Arabidopsis Protoplasts to Study Cellular Responses to Environmental Stress. in Environmental Responses in Plants: Methods and Protocols (ed. Duque, P.) 247–269 (Springer, 2016). doi:10.1007/978-1-4939-3356-3_20.

28. Asai, T. et al. Fumonisin B1-induced cell death in arabidopsis protoplasts requires jasmonate-, ethylene-, and salicylate-dependent signaling pathways. Plant Cell 12, 1823–1836 (2000).

29. Sheen, J. Signal Transduction in Maize and Arabidopsis Mesophyll Protoplasts. Plant Physiol. 127, 1466–1475 (2001).

30. Miao, Y. & Jiang, L. Transient expression of fluorescent fusion proteins in protoplasts of suspension cultured cells. Nature Protocols 2, 2348–2353 (2007).

31. Yoo, S. D., Cho, Y. H. & Sheen, J. Arabidopsis mesophyll protoplasts: a versatile cell system for transient gene expression analysis. Nature Protocols 2, 1565–1572 (2007).

32. Wu, F. H. et al. Tape-Arabidopsis Sandwich - a simpler Arabidopsis protoplast isolation method. Plant Methods 5, (2009).

33. Martinho, C. et al. Dissection of miRNA Pathways Using Arabidopsis Mesophyll Protoplasts. Molecular Plant 8, 261–275 (2015).

34. Nanjareddy, K., Arthikala, M.-K., Blanco, L., Arellano, E. S. & Lara, M. Protoplast isolation, transient transformation of leaf mesophyll protoplasts and improved Agrobacterium-mediated leaf disc infiltration of Phaseolus vulgaris: tools for rapid gene expression analysis. BMC Biotechnol 16, 53 (2016).

35. Kim, J.-Y. et al. Distinct identities of leaf phloem cells revealed by single cell transcriptomics. The Plant Cell (2021) doi:10.1093/plcell/koaa060.

36. Samodelov, S. L. et al. StrigoQuant: A genetically encoded biosensor for quantifying strigolactone activity and specificity. Science Advances 2, e1601266 (2016).

37. Aguilar-Martinez, J. A., Poza-Carrion, C. & Cubas, P. Arabidopsis BRANCHED1 acts as an integrator of branching signals within axillary buds. Plant Cell 19, 458–472 (2007).

38. Finlayson, S. A. Arabidopsis Teosinte Branched1-like 1 regulates axillary bud outgrowth and is homologous to monocot Teosinte Branched1. Plant Cell Physiol. 48, 667–677 (2007).

39. González-Grandío, E. et al. Abscisic acid signaling is controlled by a <em>BRANCHED1/HD-ZIP I</em> cascade in <em>Arabidopsis</em> axillary buds. Proc Natl Acad Sci USA 114, E245 (2017).

40. Schulz, M. H. et al. DREM 2.0: Improved reconstruction of dynamic regulatory networks from time-series expression data. BMC Systems Biology 6, 104 (2012).

41. Sarnowska, E. et al. The Role of SWI/SNF Chromatin Remodeling Complexes in Hormone Crosstalk. Trends in Plant Science 21, 594–608 (2016).

42. Bargmann, B. O. R. et al. TARGET: A Transient Transformation System for Genome-Wide Transcription Factor Target Discovery. Molecular Plant 6, 978–980 (2013).

43. Buenrostro, J. D., Giresi, P. G., Zaba, L. C., Chang, H. Y. & Greenleaf, W. J. Transposition of native chromatin for fast and sensitive epigenomic profiling of open chromatin, DNA-binding proteins and nucleosome position. Nature Methods 10, 1213–+ (2013).

44. Yu, G., Wang, L.-G. & He, Q.-Y. ChIPseeker: an R/Bioconductor package for ChIP peak annotation, comparison and visualization. Bioinformatics 31, 2382–2383 (2015).

45. O’Malley, R. C. et al. Cistrome and Epicistrome Features Shape the Regulatory DNA Landscape. Cell 165, 1280–1292 (2016).

46. Tsompana, M. & Buck, M. J. Chromatin accessibility: a window into the genome. Epigenetics & Chromatin 7, (2014).

47. Lu, Z., Ricci, W. A., Schmitz, R. J. & Zhang, X. Identification of cis-regulatory elements by chromatin structure. Current Opinion in Plant Biology 42, 90–94 (2018).

48. Shu, J. et al. Genome-wide occupancy of Arabidopsis SWI/SNF chromatin remodeler SPLAYED provides insights into its interplay with its close homolog BRAHMA and Polycomb proteins. Plant J (2021) doi:10.1111/tpj.15159.

49. Guo, J. et al. Comprehensive characterization of three classes of Arabidopsis SWI/SNF chromatin remodelling complexes. Nat. Plants 1–17 (2022) doi:10.1038/s41477-022-01282-z.

50. Brewer, P. B., Koltai, H. & Beveridge, C. A. Diverse Roles of Strigolactones in Plant Development. Molecular Plant 6, 18–28 (2013).

51. Kerr, S. C. et al. Hormonal regulation of the BRC1-dependent strigolactone transcriptome involved in shoot branching responses. http://biorxiv.org/lookup/doi/10.1101/2020.03.19.999581 (2020) doi:10.1101/2020.03.19.999581.

52. Xie, M. et al. A B-ARR-mediated cytokinin transcriptional network directs hormone cross-regulation and shoot development. Nat Commun 9, 1604 (2018).

53. Lewis, D. R. et al. A Kinetic Analysis of the Auxin Transcriptome Reveals Cell Wall Remodeling Proteins That Modulate Lateral Root Development in Arabidopsis[W][OPEN]. Plant Cell 25, 3329–3346 (2013).

54. Liu, Y. et al. Genome-wide mapping of DNase I hypersensitive sites reveals chromatin accessibility changes in Arabidopsis euchromatin and heterochromatin regions under extended darkness. Scientific Reports 7, 4093 (2017).

55. Wang, P. et al. Chromatin accessibility and translational landscapes of tea plants under chilling stress. Horticulture Research 8, 1–15 (2021).

56. Han, S.-K. et al. The SWI2/SNF2 Chromatin Remodeling ATPase BRAHMA Represses Abscisic Acid Responses in the Absence of the Stress Stimulus in *Arabidopsis*. Plant Cell 24, 4892–4906 (2012).

57. Paro, R., Grossniklaus, U., Santoro, R. & Wutz, A. Biology of Chromatin. in Introduction to Epigenetics (eds. Paro, R., Grossniklaus, U., Santoro, R. & Wutz, A.) 1–28 (Springer International Publishing, 2021). doi:10.1007/978-3-030-68670-3_1.

58. Ma, H. et al. A D53 repression motif induces oligomerization of TOPLESS corepressors and promotes assembly of a corepressor-nucleosome complex. Sci. Adv. 3, e1601217 (2017).

59. Wagner, D. & Meyerowitz, E. M. SPLAYED, a Novel SWI/SNF ATPase Homolog, Controls Reproductive Development in Arabidopsis. Current Biology 12, 85–94 (2002).

60. Farrona, S., Hurtado, L., Bowman, J. L. & Reyes, J. C. The Arabidopsis thaliana SNF2 homolog AtBRM controls shoot development and flowering. Development 131, 4965–4975 (2004).

61. Tang, X. et al. The Arabidopsis BRAHMA Chromatin-Remodeling ATPase Is Involved in Repression of Seed Maturation Genes in Leaves. Plant Physiol 147, 1143–1157 (2008).

62. Hurtado, L., Farrona, S. & Reyes, J. C. The putative SWI/SNF complex subunit BRAHMA activates flower homeotic genes in Arabidopsisthaliana. Plant Mol Biol 62, 291–304 (2006).

63. Bajic, M., Maher, K. A. & Deal, R. B. Identification of Open Chromatin Regions in Plant Genomes Using ATAC-Seq. in Plant Chromatin Dynamics (eds. Bemer, M. & Baroux, C.) vol. 1675 183–201 (Springer New York, 2018).

64. Babraham Bioinformatics - FastQC A Quality Control tool for High Throughput Sequence Data. http://www.bioinformatics.babraham.ac.uk/projects/fastqc/ (2019).

65. Bolger, A. M., Lohse, M. & Usadel, B. Trimmomatic: a flexible trimmer for Illumina sequence data. Bioinformatics 30, 2114–2120 (2014).

66. Patro, R., Duggal, G., Love, M. I., Irizarry, R. A. & Kingsford, C. Salmon provides fast and bias-aware quantification of transcript expression. Nature Methods 14, 417–419 (2017).

67. Love, M. I., Huber, W. & Anders, S. Moderated estimation of fold change and dispersion for RNA-seq data with DESeq2. Genome Biology 15, (2014).

68. Conway, J. R., Lex, A. & Gehlenborg, N. UpSetR: an R package for the visualization of intersecting sets and their properties. Bioinformatics 33, 2938–2940 (2017).

69. Bindea, G. et al. ClueGO: a Cytoscape plug-in to decipher functionally grouped gene ontology and pathway annotation networks. Bioinformatics 25, 1091–1093 (2009).

70. Bourbousse, C., Vegesna, N. & Law, J. A. SOG1 activator and MYB3R repressors regulate a complex DNA damage network in *Arabidopsis*. Proc Natl Acad Sci USA 115, E12453–E12462 (2018).

71. Langmead, B. & Salzberg, S. L. Fast gapped-read alignment with Bowtie 2. Nat Methods 9, 357–359 (2012).

72. Ramírez, F., Dündar, F., Diehl, S., Grüning, B. A. & Manke, T. deepTools: a flexible platform for exploring deep-sequencing data. Nucleic Acids Res 42, W187–W191 (2014).

73. Zhang, Y. et al. Model-based Analysis of ChIP-Seq (MACS). Genome Biology 9, R137 (2008).

74. Stark, R. & Brown, G. DiffBind: Differential binding analysis of ChIP-Seq peak data. 33.

75. Bailey, T. L. STREME: Accurate and versatile sequence motif discovery. Bioinformatics (2021) doi:10.1093/bioinformatics/btab203.

76. Gupta, S., Stamatoyannopoulos, J. A., Bailey, T. L. & Noble, W. S. Quantifying similarity between motifs. Genome Biology 8, R24 (2007).

77. Grant, C. E., Bailey, T. L. & Noble, W. S. FIMO: scanning for occurrences of a given motif. Bioinformatics 27, 1017–1018 (2011).

